# Deciphering the dynamic proteome of multiciliated cells

**DOI:** 10.1101/2025.06.10.658933

**Authors:** Camille Boutin, Olivier Rosnet, Marine Touret, Stéphane Audebert, Luc Camoin, Salomé Dussert, Nicolas Brouilly, Virginie Thomé, Denis Fortun, Jean-Paul Borg, Laurent Kodjabachian

## Abstract

Multiciliated cells (MCCs) are essential for generating directional fluid flow across specialized epithelia in various vertebrate organs. MCC differentiation involves a unique, tightly regulated program characterized by massive centriole amplification, independently of DNA replication. Although much is known about the transcriptional control of MCC development, insights into proteome dynamics have been limited due to the lack of suitable models. In this study, we report the generation of a stable inducible MCC line, derived from *Xenopus laevis* A6 kidney epithelial cells. Upon induction of the MCC master regulator Multicilin (MCI), most A6-MCI cells synchronously differentiate into mature MCCs in 48 hours. Using this novel resource, custom antibodies and super-resolution imaging, we characterized *Xenopus* deuterosomes, the platforms that allow massive centriole synthesis in vertebrate MCCs. We performed a detailed proteomic profiling throughout MCC differentiation, and uncovered previously uncharacterized regulators. Notably, we highlight a critical role for CDK7 in MCC differentiation in both *Xenopus* and human systems. Our work provides a valuable resource for mechanistic studies of MCC biology and opens avenues to identify novel therapeutic targets for motile ciliopathies.

## Introduction

Multiciliated cells (MCCs) are widely present in the evolutionary tree and support major functions by generating physiological fluid flow at the surface of specialized epithelia through beating of hundreds of cilia. In humans, MCCs are essential for the circulation of the cerebro-spinal fluid in the central nervous system, in transportation of gametes, and in evacuation of soiled mucus from upper airways (Spassky and Meunier, 2017). Mutations that impair MCC differentiation or function cause severe multi-symptomatic diseases such as Reduced Generation of Multiple Cilia (RGMC) and Primary Ciliary Dyskinesia (PCD) collectively referred to as motile ciliopathies (Wallmeier et al., 2020).

MCC differentiation is a highly regulated multistep process that can be described as an alternative cell cycle in which centriole amplification occurs uncoupled from DNA synthesis. MCCs amplify almost simultaneously dozens to hundreds of centrioles using centriole-dependent and deuterosome-dependent pathways (Spassky and Meunier, 2017). Following amplification in the cytoplasm, centrioles move to the apical cell surface, gain appendages to allow linkage to the cortical cytoskeleton, and nucleate the formation of motile cilia (Boutin and Kodjabachian, 2019). Beyond similarities in the cellular mechanisms at play, it has become clear that, at various levels, regulatory pathways of MCC differentiation rely on intricate coordination between canonical cell cycle regulators and MCC-specific regulators. At the transcriptional level, two factors related to the S-phase regulator Geminin: Multicilin (MCI, encoded by MCIDAS gene) and GemC, in complex with members of the E2F family of cell cycle transcriptional regulators, regulate the expression of a large body of effectors required for MCC differentiation (Kim et al., 2018; Lewis et al., 2023; Ma et al., 2014; Stubbs et al., 2012). MCI was reported to activate the expression of the transcription factor Myb, well known for its S-phase promoting activity in a variety of progenitor cells. In MCCs, Myb is required for multiple centriole synthesis and, together with MCIDAS, for the activation of FoxJ1, a critical transcriptional regulator of motile ciliogenesis (Pan et al., 2014; Quigley and Kintner, 2017; Tan et al., 2013). Most of the centrioles in MCCs are produced by specialized structures called deuterosomes (Anderson and Brenner, 1971; Brenner, 1969; Kalnins and Porter, 1969; Sorokin, 1968; Steinman, 1968). The core of the deuterosome is composed of the MCC-specific protein Deup1, a paralog of Cep63 (Zhao et al., 2013). Additional proteins such as Pericentrin and γ-Tubulin compose the peri-deuterosomal material (Revinski et al., 2018) and procentriole nucleation around deuterosomes involve many of the key players of the centriole duplication pathway, including PLK4, CEP152 and SAS6 (Al Jord et al., 2014; Klos Dehring et al., 2013; Zhao et al., 2013). At the end of the process, CDC20B – an MCC-specific protein related to the cell cycle protein CDC20 – triggers a Separase-dependent proteolytic event required for centriole disengagement from deuterosomes (Revinski et al., 2018).

The progression of MCCs through phases of differentiation is dependent on the cell cycle regulators Cyclin Dependent Kinases (CDK) (Al Jord et al., 2017; Choksi et al., 2024; Serizay et al., 2025; Vladar et al., 2018). The mitotic oscillator, comprising CDK1 and APC/C, controls centriole synthesis and disengagement (Al Jord et al., 2017). CDK2 activity regulates both early phases of MCC differentiation and ciliogenesis (Vladar et al., 2018). CDK4 and CDK6, which are regulators of G1-to-S progression, are required to initiate MCC differentiation (Choksi et al., 2024).

Most of our knowledge on the molecular control of MCC differentiation is derived from bulk or single cell transcriptomic approaches in *Xenopus*, mouse and human (Ma et al., 2014; Redman et al., 2024; Revinski et al., 2018; Serizay et al., 2025). However, information on protein expression dynamics is critically missing mainly due to the absence of model systems suitable for proteomic approaches.

In this study, we report the development of an inducible multiciliated cell line (A6-MCI). Building on the fact that MCI is necessary and sufficient to induce differentiation into MCCs of epithelial cells from *Xenopus* epidermis or mouse ependyma (Kyrousi et al., 2015; Stubbs et al., 2012), we demonstrate that the stable transfection of an inducible form of MCI is sufficient to drive differentiation into MCCs of A6, a cell line derived from the kidney of the South African clawed toad, *Xenopus laevis* (Rafferty, 1969). The A6-MCI line has a high differentiation rate and is very homogeneous, making it an ideal tool for advanced microscopy and proteomics approaches. Using this unique resource, we further characterized the organization of *Xenopus* MCC deuterosomes, revealing similar as well as unique features, when compared to mammalian MCCs. We assembled the proteome of MCCs at different stages of differentiation, which revealed several uncharacterized regulators that may contribute to key functions in MCC biology. Highlighting the value of this unique dataset, we uncovered the importance of CDK7 activity for the differentiation of *Xenopus* and human MCCs.

## Results

### MCI expression is sufficient to drive differentiation of A6 cells into MCCs

In an attempt to generate a MCC line, we tested whether forced MCIDAS expression alone could trigger MCC differentiation in established vertebrate cell lines. We first transfected RPE1 and NIH3T3 cells with mouse MCIDAS, and analyzed their differentiation using centrioles and cilia markers. We observed that neither cell lines were responsive to MCIDAS expression alone (Fig.1A, B). Next, we transfected A6 *Xenopus laevis* cells with *Xenopus laevis* MCIDAS fused to the ligand binding domain of the human glucocorticoid receptor (xMCIDAS-hGR). This inducible form of MCIDAS is retained in the cytoplasm until dexamethasone is added to the culture medium (Stubbs et al., 2012). Observation of the culture 24h after dexamethasone treatment revealed amplified centrioles and multiple cilia showing that expression of MCIDAS alone was sufficient to induce A6 cells into MCCs (Fig.1C, D). In the next step, we stably transfected A6 cells with xMCIDAS-hGR. After selection and cloning by limit dilution, we obtained the A6-MCI cell line. 72 hours post induction (hpi) with dexamethasone, A6-MCI MCCs were fully differentiated with multiple centrioles and cilia at their apical surface (Fig.1E, F). To define the best induction protocol, we compared differentiation rates in the cultures at 72hpi with increasing concentrations of dexamethasone applied for either 8 or 24h. In the case of a short induction (8h), the number of cells with multiple centrioles increased proportionally to the amount of dexamethasone added, ranging from 20% at 1nM to above 60% at 1μM (Suppl. Fig.1A, C). In the case of a prolonged induction (24h), differentiation levels were more comparable (over 50% at 1nM, up to 70% at 1μM) (Suppl. Fig.1A, D). Similar results were observed at 1week post induction, suggesting that the maximum of differentiation is reached by 72h (Suppl. Fig.1A, E). Of note, no signs of differentiation were observed in non-induced A6-MCI cultured for 72h or 1 week (Suppl. Fig.1B) suggesting that dexamethasone control of induction is robust and that there is no leakage in the system. No major differences were observed in the number of centrioles generated by individual MCCs with the different dexamethasone concentrations and those numbers were comparable with those of native *Xenopus* epidermal MCCs (Fig.1G). Finally, we observed that in all conditions, over 80% of cells that undergo centriole amplification also make multicilia (Fig.1H). Following these analyses, induction for 24 hours with 100 nM dexamethasone was selected as the routine condition.

**Figure 1.**
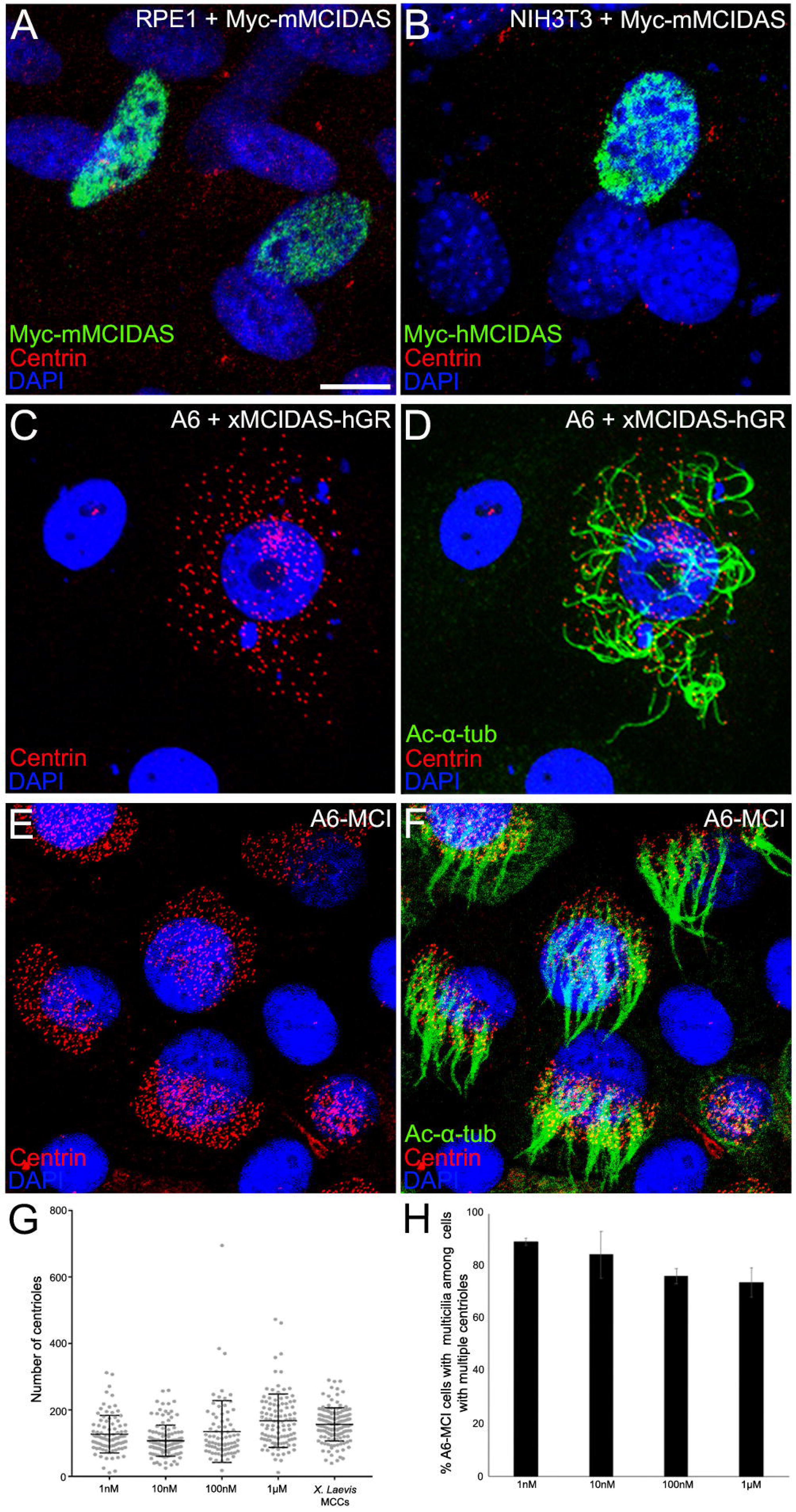
MCI expression in A6 cells drives differentiation in MCCs. (A-D) Confocal pictures of RPE1 (A), NIH3T3 (B) and A6 (C, D) cells transiently transfected with Myc-mMCIDAS or XMCIDAS-hGR and immunostained for indicated markers. (E-F) Confocal picture of A6-MCI cells stained for indicated markers. (G) Quantification of centriole number per cell in A6-MCI induced with increasing concentration of dexamethasone and in *Xenopus laevis* epidermal MCCs. n=91 cells (1nM), 124 cells (10nM), 80 cells (100nM), 103 cells (1μM) and 155 cells (*X. laevis*). Data collected from one experiment. (H) Quantification of percentage of cells with multicilia among cells with multiple centrioles in A6-MCI cells induced with increasing concentrations of Dexamethasone. n=936 (1nM), 1277 (10nM), 1432 (100nM) and 1282 (1μM) cells. Data collected from two experiments. Scale bar: 10μm (A-D), 13μm (E-F).

Altogether, these results demonstrate that the stable expression of MCIDAS is alone sufficient to induce a rapid, robust and reproducible differentiation of A6 cells into MCCs.

### A6-MCI differentiation is synchronous and recapitulates the main features of MCCs

To further characterize the A6-MCI line, we performed a time course analysis to define the main steps of their differentiation. As MCC differentiation starts with centriole synthesis, we aimed at analyzing the expression of Deup1, which constitutes the core of vertebrate deuterosomes, the platforms where massive centriole assembly occurs. For this, we raised and characterized a rabbit polyclonal antibody against *Xenopus* Deup1, which proved to give specific signals in both Western blotting and immunofluorescence (Fig.2A-C and Suppl. Fig.2A, B). In Western blots, Deup1 was indeed the first MCC-specific protein detected. It appeared as soon as 8hpi, increased to reach a plateau at 24hpi before declining until being only weakly detected one week after induction. As expected, the increase in expression of the centriole marker Centrin was slightly delayed compared to Deup1. While low basal expression was detected in NI and at 8hpi, Centrin strongly increased from 12hpi up to 48hpi and then remained stable. Finally, the acetylated form of alpha-Tubulin (Ac-α-Tubulin), a cilia component, increased from baseline levels in non-induced cells to reach maximum levels at 72hpi, and remained stable up to one week post induction (Suppl. Fig.2C). Next, we performed immunostainings to characterize the state of MCC differentiation at different times post-induction. At 8hpi, although the Western blot data showed that the differentiation program was engaged, induced and non-induced cells appeared similar with respect to deuterosome (Deup1) and centriole (Centrin) markers (Fig.2A, B, J). At 16 hpi, over 50% of induced cells were engaged in the phase of centriole amplification as shown by accumulation of structures positive for Deup1, γ-Tubulin, Pericentrin and Centrin, corresponding to deuterosomes bearing pro-centrioles (Fig.2C, J, K; Suppl. Fig. 3). At 24 hpi, centriole amplification was almost completed, as shown by the decrease in the number of cells with deuterosomes and increase in the number of cells with released centrioles (Fig.2D-F, J, K). In most cells, individual centrioles were transforming into basal bodies, as shown by the presence of γ-Tubulin and Cep164 - respective markers of the basal foot and distal appendages - juxtaposed to Centrin (Fig.2 E, F). At 24hpi, cilia were not grown however IFT88 started to accumulate at basal bodies in a subset of cells (Suppl. Fig.3). At 48hpi, a majority of cells presented cilia at their apical surface (Fig.2G). Accordingly, ciliary pools of IFT88 were observed indicating active transport into cilia (Fig.2H). In addition, Ruvb2-positive Dynaps, which have been described as condensates enabling the association of dynein arms prior to their import into cilia (Huizar et al., 2018), were observed in most of the cells (Fig.2I). Finally, at 72 hpi, deuterosomes and Dynaps were no longer observed and most cells presented mature centrioles and were ciliated (Suppl. Fig.3). Next, we used high-resolution imaging to better assess the structures of centrioles and cilia in A6-MCI MCCs. Ultrastructure Expansion Microscopy (UExM) revealed the expected distal position of Centrin within the space delineated by centriole walls, marked by Ac-α-Tub (Fig. 3A). UExM also confirmed the nine-fold distribution of Cep164-positive distal appendages, as well as the asymmetric position of the γ-Tubulin-positive basal foot (Fig.3B-D). We further analyzed A6-MCI centrioles using Transmission Electron Microscopy (TEM). As classically observed in MCCs, A6-MCI centrioles presented nine microtubule triplets and were decorated with a basal foot linked to cytoplasmic microtubules, nine distal appendages and rootlets (fan-shaped or long and slim, like in *Xenopus* epidermal MCCs) (Fig.3E-H). Altogether, these results show that over 70% A6-MCI induced cells recapitulate the main steps of MCC differentiation in a precisely timed manner.

**Figure 2.**
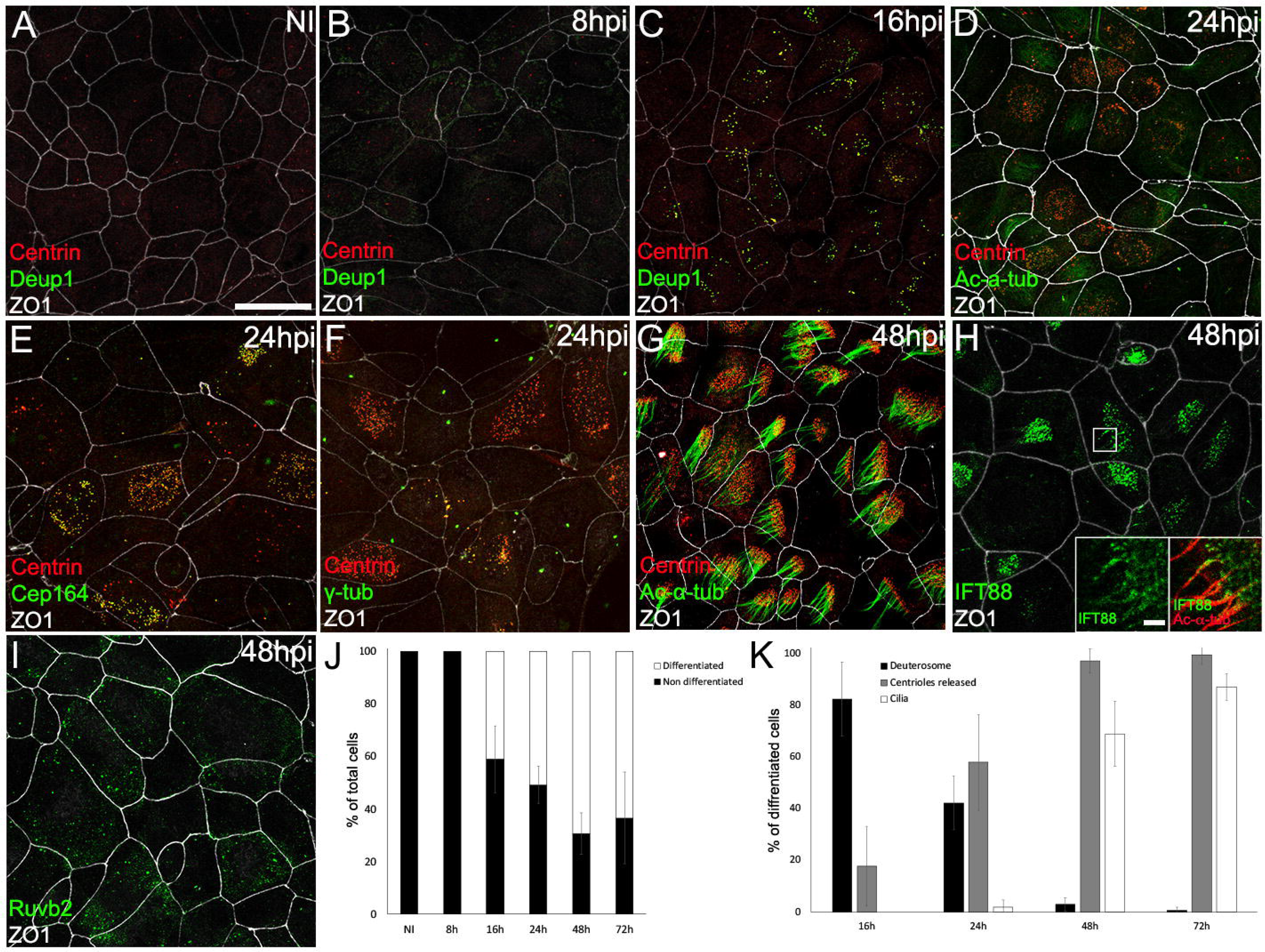
A6-MCI cells recapitulates main differentiation steps of MCCs. (A-I) Confocal pictures of non-induced (NI, A) or induced A6-MCI cells (B-I) at indicated hours post induction (hpi) stained for indicated markers. (J) Bar graph quantification of non-differentiated cells and differentiated cells at indicated time post induction. For stages NI, 8h, and 16, cell differentiation was evaluated by the presence of Deup1 and Centrin labelling. For stages 24h, 48h, and 72h, cell differentiation was evaluated by the presence of Centrin labelling. n= 414 (NI), 456 (8h), 293 (16h), 288 (24h), 478 (48h) and 373 (72h) cells. Data were collected from three independent experiments. (K) Bar graph showing the distribution of differentiated cells at 16hpi, 24hpi and 72hpi in deuterosome, centriole released and ciliated categories evaluated by Deup1 and Centrin, Centrin and Ac-α-Tubulin staining respectively. n=115 (16h), 288 (24h), 148 (48h) and 330 (72h) cells. Data were collected from three independent experiments. Scale bar: 30μm (A-I); 2.5μm (G, inset).

**Figure 3.**
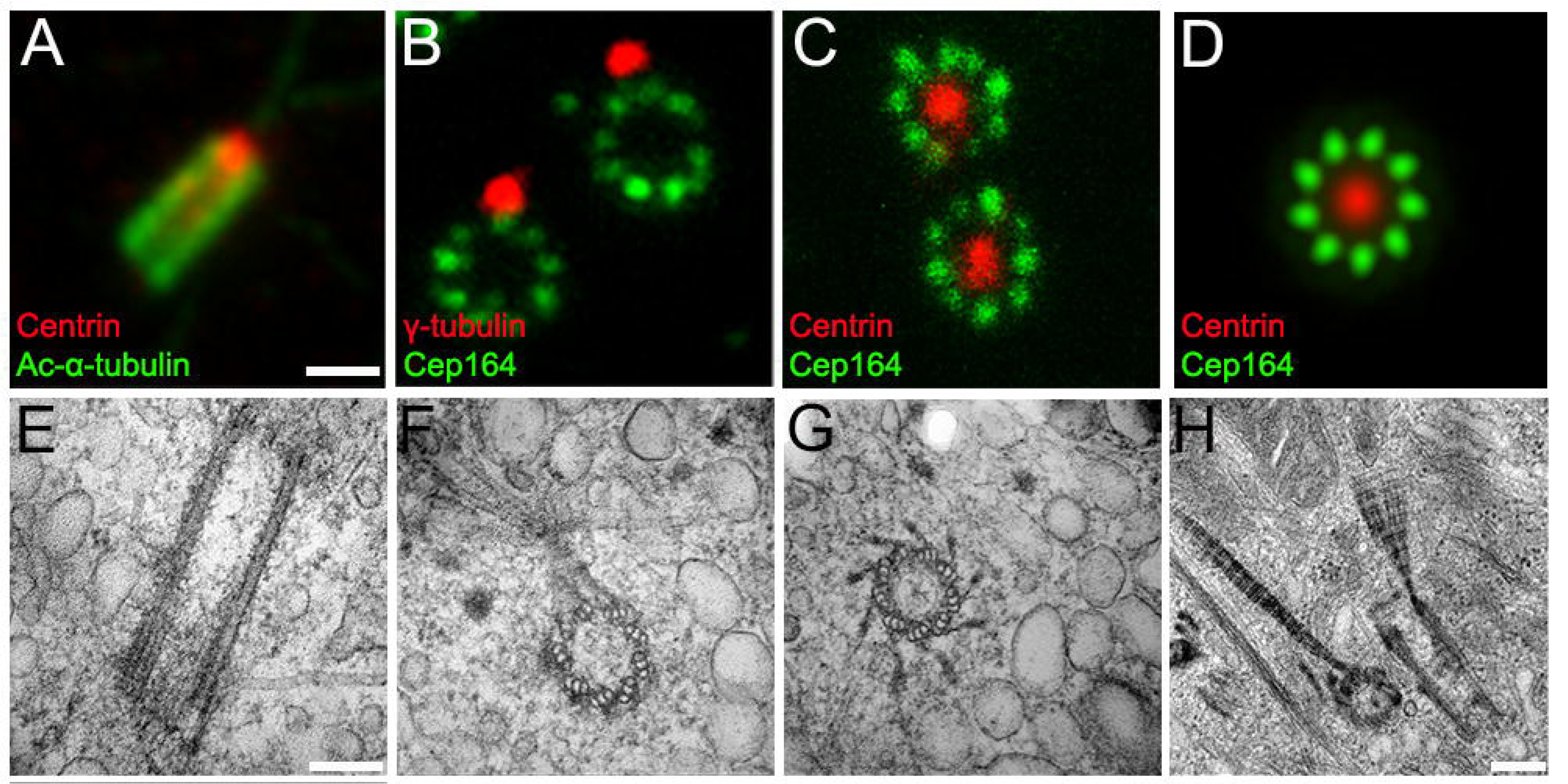
Centrioles in A6-MCI cells. (A-C) Confocal pictures of basal bodies of expanded differentiated A6-MCI cells stained for Acetylated-α-tubulin (microtubule) and centrin (centriole) (A), γ-tubulin (basal foot) and Cep164 (distal appendages) (B) or Cep164 (distal appendages) and centrin (centriole) (C). (D) Average map distribution of microtubules, centrioles, basal foot and distal appendages reconstructed from 13 centrioles. (E-H) TEM pictures from differentiated A6-MCI cells. (E) Lateral view of centriole. (F) Top view of centriole displaying 9 triplet microtubules centriolar wall and basal foot nucleating microtubules. (G) Top view of centriole displaying 9 triplet microtubules centriolar wall and distal appendages. (H) Top and lateral views of basal bodies showing rootlet. Scale bar: 250nm (A-C); 150nm (E-G); 200nm (H)

### *Xenopus* deuterosomes exhibit a distinctive organization

In vertebrate MCCs, deuterosomes support the production of the major part of the total number of centrioles (Spassky and Meunier, 2017). In the *Xenopus* embryonic epidermis, cells at the deuterosomal stage are localized in the inner layer and represent about 10% of the cells present in the tissue, which makes high-resolution imaging of the sub-micrometric deuterosome organelle challenging (Boutin and Kodjabachian, 2019). Thus, we decided to use the A6-MCI model to investigate the molecular and architectural organization of *Xenopus* deuterosomes. We first analyzed with confocal microscopy the respective distribution of Deup1 and Centrin signals in deuterosomal cells. Consistent with our previously published observations (Revinski et al., 2018), the deuterosomes of both A6-MCI MCCs and *Xenopu*s epidermal MCCs showed irregular shapes and sizes (Fig.4A, B). We then analyzed the distribution of γ-Tubulin and Pericentrin, two proteins that we had previously identified as being associated with the deuterosome of *Xenopus* epidermal and mouse ependymal MCCs (Revinski et al., 2018). Confocal immunofluorescence confirmed that those proteins localized to A6-MCI deuterosomes associated to Centrin-positive centrioles (Fig.4C, D). Together, these results suggest that A6-MCI deuterosomes are similar to that observed in natural *Xenopus* MCCs. Interestingly, although the shapes of deuterosomes differ between mouse ependymal and *Xenopus* MCCs, STED super-resolution microscopy analysis on A6-MCI MCCs revealed that the relative position of markers are conserved, with Deup1 being central, γ-Tubulin and Pericentrin being more peripheral in that order (Suppl. Fig.4A; (Revinski et al., 2018)).

**Figure 4.**
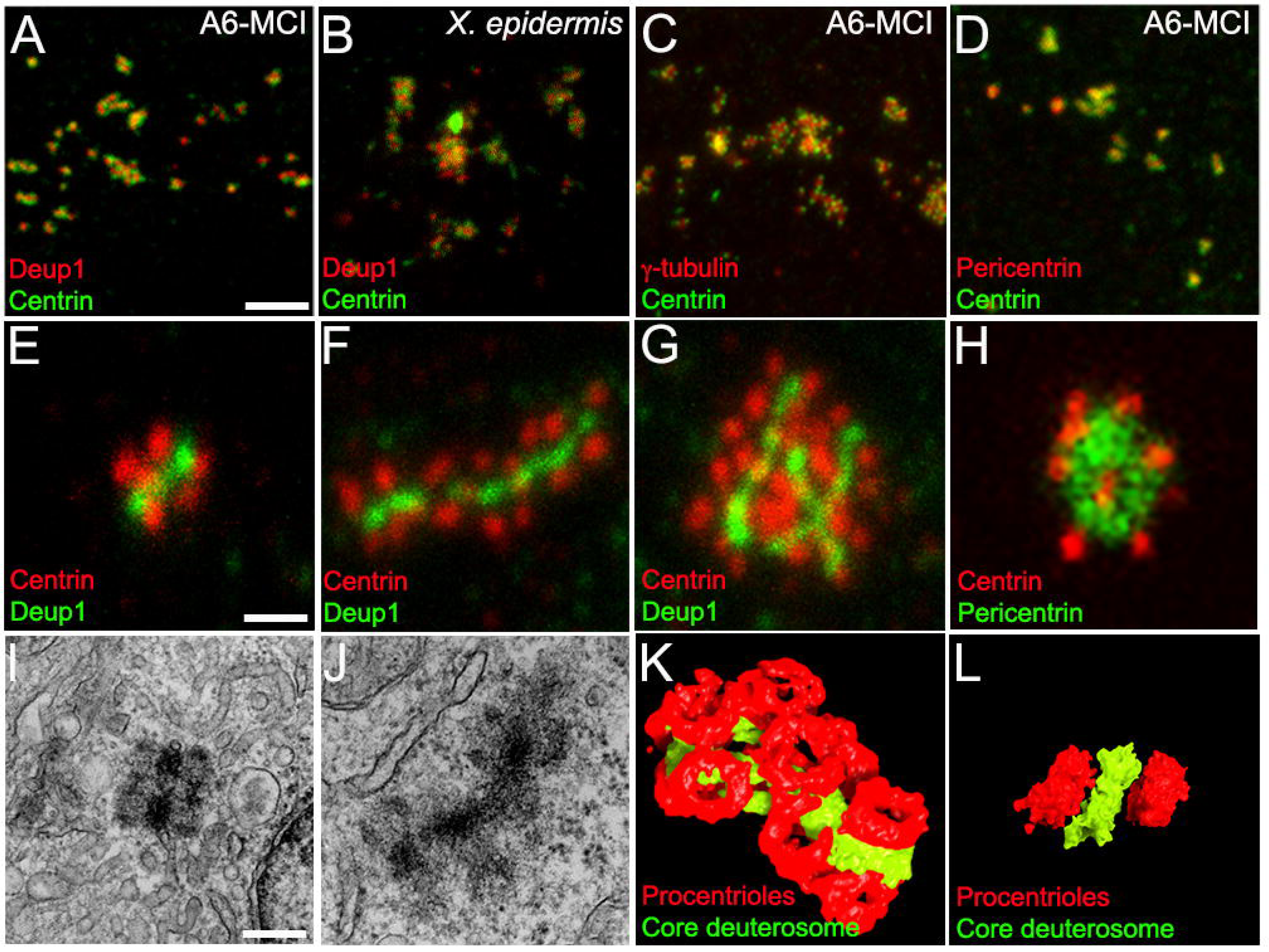
Deuterosomes organization in A6-MCI cells. (A-D) Confocal pictures of 16hpi A6-MCI (A, C, D) or *Xenopus* epidermis (B) MCCs stained with indicated deuterosome (Deup1, Pericentrin, γ-tubulin) or centriole (centrin) markers. (E-G) Expansion microscopy picture of short (E), long (F) and ramified (G) A6-MCI deuterosomes stained for Deup1 and Centrin. (H) STED picture of globular A6-MCI deuterosome stained with pericentrin and centrin. (I-J) TEM picture of A6-MCI deuterosomes. (K-L) Segmentation obtained from Tomogram acquisition of ramified (K) and short (L) A6-MCI deuterosomes. Scale bar: 2.5 μm (A-D); 280nm (E-H); 225nm (I-J).

Next, we used fourfold magnification UExM to further analyze the shape of deuterosomes in A6-MCI cells. Three major shapes were observed: short deuterosomes with a central Deup1-positive core of approximately 300 to 400 nm bearing 2 to 6 procentrioles (Fig.4E); long deuterosomes, which presented a linear core between 1.75μm and 3.15 μm long and carried 18 to 28 procentrioles (Fig.4F); and finally, larger deuterosomal structures with Deup1-positive segments ramified (Fig.4G). Deuterosomes with globular cores were occasionally observed (Fig.4H). Upon closer observation, long deuterosomal entities appeared to be formed of juxtaposed spheroid Deup1-positive units. To extend this analysis, we processed A6-MCI cells for serial TEM. Using this approach, we could observe small deuterosomes loaded with few procentrioles, restricted to one section (thickness= 70nm), as well as long deuterosomes spanning several consecutive sections (Fig.4I, J; Suppl. Fig.4B). However, the serial TEM approach did not allow us to precisely reconstruct the 3D structure of deuterosomes spanning more than one section. Therefore, we turned to serial tomography, which allows the reconstruction of EM volumes at high and isotropic resolution. With this approach, we were able to reconstruct in 3D, small, long and ramified structures thereby confirming the existence of various deuterosomal architecture (Fig.4K, L; Suppl. Movie 1-3). Importantly, ultrastructure analysis with TEM and tomography confirmed the “string of pearls” organization of deuterosomal platforms, revealed by UExM. Regardless of the size or general shape of deuterosomal platforms, most of them appeared as a series of electron dense spheroid (86±14nm diameter) connected by thinner material, each bearing two to three procentrioles. Altogether, these results show that A6-MCI cells contain deuterosomes similar to those of *Xenopus* epidermal MCCs. High-resolution imaging also suggests that deuterosomes of *Xenopu*s MCCs are composed of small spherical units that can be linked to generate supra-structures of variable length and geometry.

### Temporal proteomic analysis of MCC differentiation

The results presented above indicate that our A6-MCI cell line recapitulate MCC differentiation with high fidelity. To further characterize this progression, we performed mass spectrometry analysis to define the list of proteins expressed in non-induced cells (NI) and at 8h, 16h, 24h, 48h and 72h post-induction (Suppl. Table 1). Principal Component Analysis (PCA) of mass spectrometry data revealed that samples were separated according to time post-induction, confirming sequential, synchronous and reproducible differentiation of A6-MCI cells (Fig.5A). We then analyzed the difference in protein expression between successive time points (Suppl. Table 1). Consistent with the absence of major signs of differentiation in immunofluorescence, little changes in protein expression were observed between NI and 8h (Fig.5B). The number of proteins up-regulated between successive time points gradually increased from 8h to 48h reflecting the progression into the different phases of differentiation of induced A6-MCI cells (Fig.5C-E). In contrast, between 48h and 72h, the differences in proteomes were less significant (Fig.5F), as expected since a majority of induced cells have completed ciliogenesis at 48hpi (Fig.2J-K). The 8h-16h transition clearly revealed the engagement into the phase of centriole synthesis, with up-regulation of Cdk1, Deup1, Plk4 and Centrin 4 (Fig.5C). At 16h-24h transition, the up-regulation of Plk1 involved centriole disengagement from deuterosome (Revinski et al., 2018), showed transitioning towards the termination of centriole production (Fig.5D). The 24h-48h transition revealed the down-regulation of Cdk1, Deup1 and Plk4, concomitantly to the up-regulation of Cdkn1a, the negative regulator of Cdk1 and Ift88 (Fig. 5E). Those changes are indeed compatible with the transition from centriole synthesis to ciliogenesis phases. To generate a more global overview of our dataset, we analyzed the top 20 gene ontology (GO) terms that describe the proteins significantly up-regulated at the 16h-24h and 24h-48h transitions. The 16h-24h transition was characterized by the presence of 4 GO terms related to centriole biogenesis, and 11 terms related to cilia assembly (Fig. 5G). In the 24h-48h transition, all centriole biogenesis GO terms were lost and 18/20 terms were related to cilia assembly and motility (Fig.5H). Overall, the data described above indicate that the A6-MCI system is amenable to proteomic analyses, as its level of synchrony is sufficient to distinguish the multiple phases of the alternative cell cycle at play in MCCs (Al Jord et al., 2017; Choksi et al., 2024; Serizay et al., 2025).

**Figure 5.**
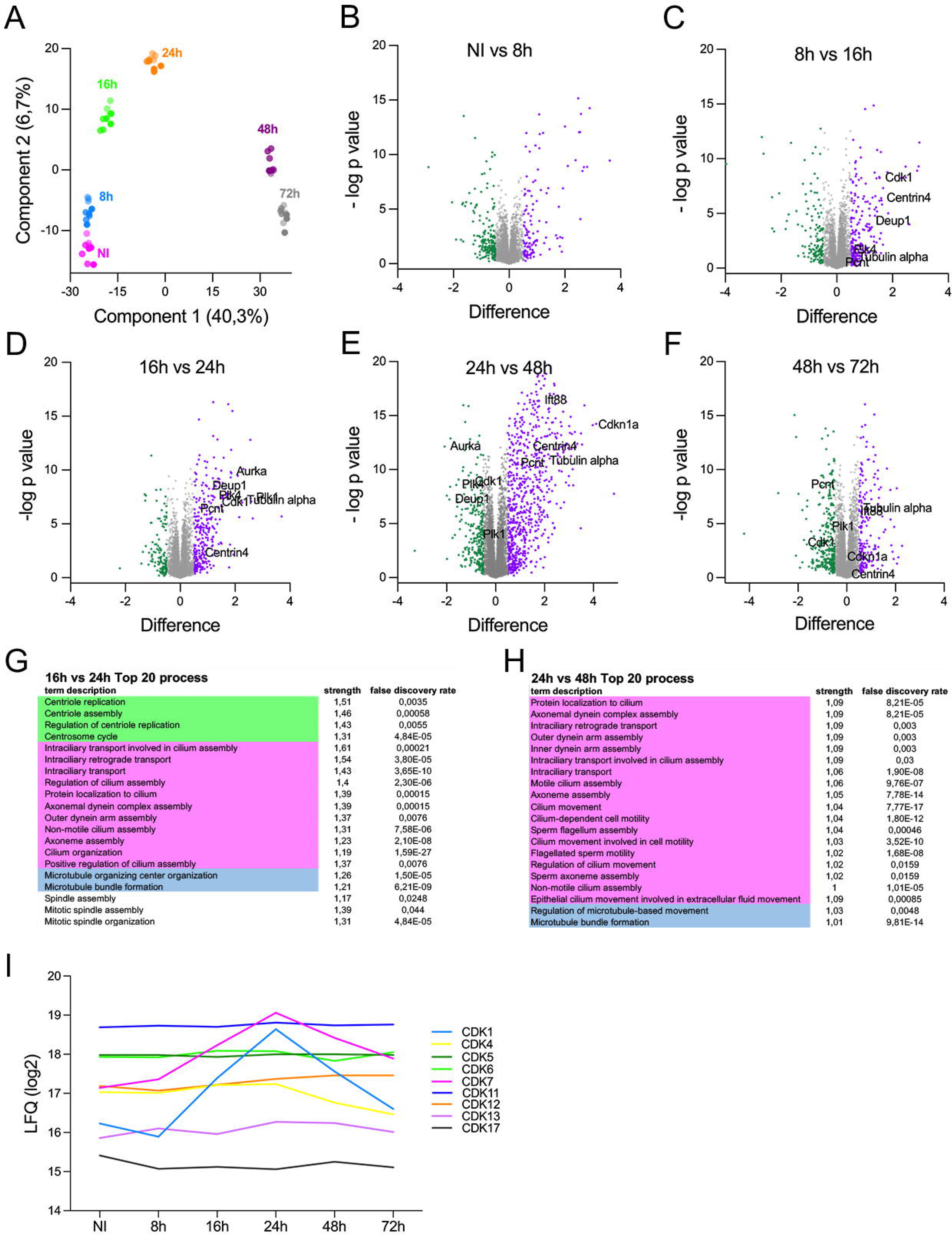
Proteomic of A6-MCI cells. (A) PCA analysis of proteomic data set. (B-F) Volcano plots of differential expression analysis between indicated time post induction. Log2 difference in expression between time point and -log p value are plotted. Up-regulated proteins with a Log2 difference >0.5 are shown in purple Down-regulated proteins with a Log2 difference <-0.5 are shown in green. Ni vs 8h 125 up 194 down; 8h vs 16h 222 up 137 down; 16h vs 24h 301up 124 down; 24h vs 48h 699 up 270 down 48h vs 72h 187 up 298 down (G-H) Top 20 GO term process associated with 16h-24h (G) and 24h-48h (H) transitions. (I) Graphical showing the profil variation of expression of all Cdks detected in the proteomic data set using the LFQ intensity.

In an attempt to identify novel regulators of the MCC alternative cycle, we focused our attention on members of the CDK family detected in the proteomes. We could detect 10 Cdks with two distinct temporal expression profiles (Fig.5I). On the one hand, Cdk1 and Cdk7 showed dynamic and parallel expression, increasing during the phase of centriole synthesis (16h-24h), before decreasing during the ciliogenesis phase (24h-72h) (Fig.5I). On the other hand, Cdk4, Cdk6, Cdk5, Cdk11, Cdk12, Cdk13 and Cdk17 showed stable expression levels between the different stages of differentiation (Fig.5I). Among the Cdk members detected in our dataset, Cdk1, Cdk4 and Cdk6 have been shown to regulate the MCC alternative cell cycle (Al Jord et al., 2017; Choksi et al., 2024; Vladar et al., 2018). Next, we analyzed whether Cdk7, the expression of which matches that of Cdk1, is important for MCC biogenesis.

### CDK7 plays a conserved role in differentiating MCCs

In *Xenopus*, Cdk7 was shown to be a direct transcriptional target of MCI (Ma et al., 2014). CDK7 in association with cyclin H (CCNH) and MAT1A drives cell cycle progression by promoting the activity of CDK1, 2, 4, and 6 (Fisher, 2005), which have all been involved in the MCC alternative cycle (Al Jord et al., 2017; Choksi et al., 2024; Vladar et al., 2018). We tested the involvement of CDK7 in the alternative cell cycle of MCCs using chemical approaches. Strikingly, the application of CDK7 inhibitors (YKL-5-124 and LDC4297) on A6-MCI cells at the time of induction almost totally prevented MCC differentiation (Fig.6A-D). To probe the validity of this result *in vivo*, we analysed epidermal MCC from *Xenopus* embryos cultured in the presence of CDK7 inhibitors. Control embryos showed homogeneously distributed, fully ciliated MCCs. In contrast, embryos treated with LDC4297 or YKL-5-124, although containing correct numbers of MCCs, showed severe differentiation defects, with most cells that failed to generate centrioles and cilia (Fig.6E-H). Next, we explored at what stage of the differentiation cascade Cdk7 was necessary. When either CDK7 inhibitor was added on A6-MCI at 8 hpi, MCC differentiation was greatly suppressed, to a rate comparable to that observed when inhibition started at induction. CDK7 inhibition from 16 hpi impaired MCC differentiation to a lower extent. Finally, when the drugs were added at 24 h or 48 h, no difference was observed compared to the control situation, suggesting that cells produced centrioles and cilia normally (Suppl. Fig.5A). Similarly, when the drug YKl-5-124 was added to *Xenopus* early gastrula (St 10) embryos, MCCs did not produce centrioles and cilia, whereas when the drug was added between the centriologenesis and ciliogenesis phases (St 20), no major defects were observed (Supl. Fig.5B). These results ruled out toxicity and confirmed the involvement of Cdk7 at an early phase, as suggested by the proteomic analysis. Published single cell transcriptomic data reveal that *CDK7*, *CCNH* and *MAT1* are also expressed in the MCC lineage of human airway epithelial cell cultures (hAEC) (Redman et al., 2024). To evaluate the importance of CDK7 in the biogenesis of human MCCs, we applied CDK7 inhibitors on air-liquid hAEC cultures at ALI day 0, when differentiation starts. Here again, both LDC4297 and YKL-5-124 nearly totally abolished MCC differentiation (Fig.6I-L). Together, these results indicate that CDK7 plays a crucial role at an early step of MCC differentiation, which is conserved from *Xenopus* to human.

**Figure 6.**
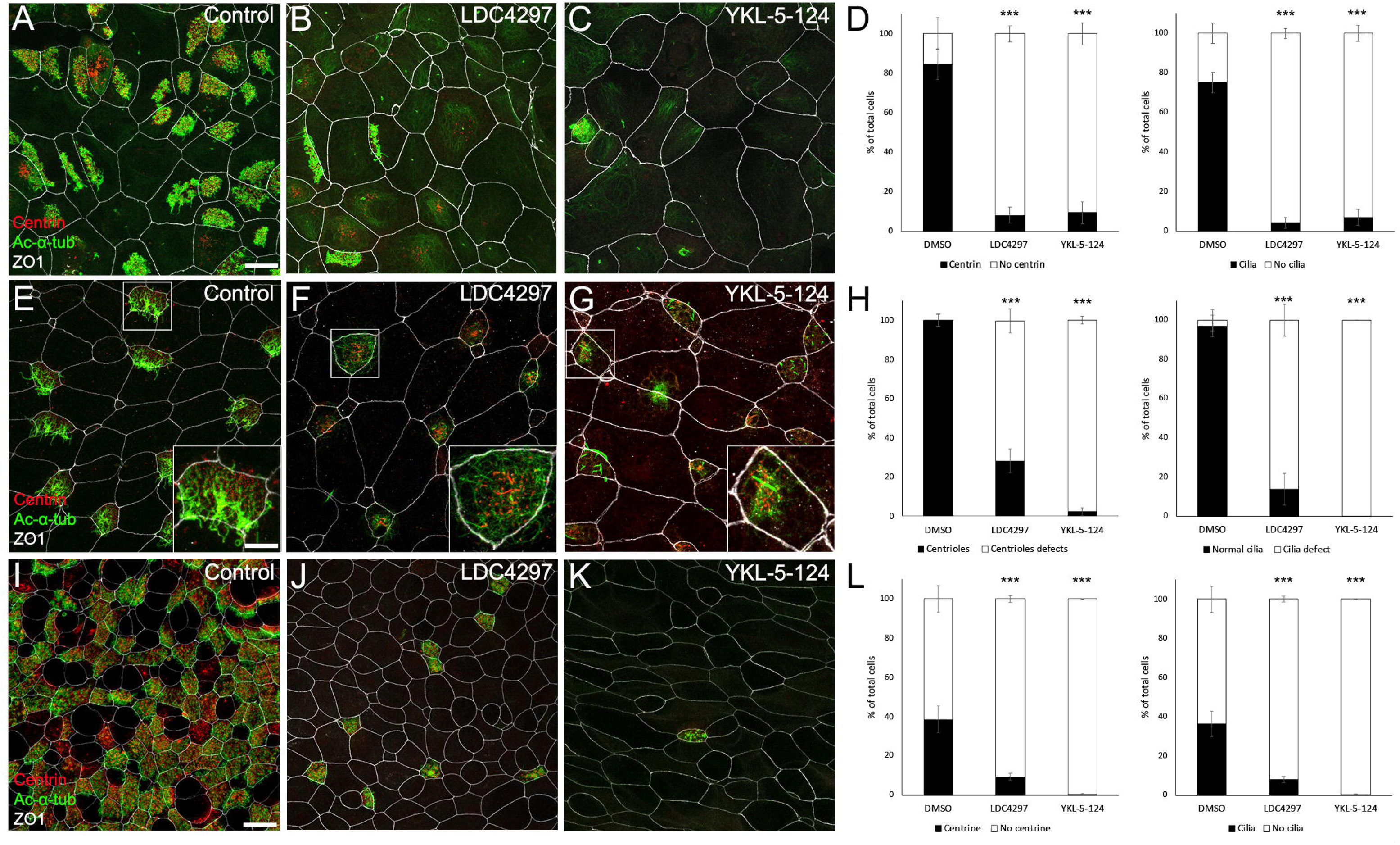
CDK7 is plays a conserved role in differentiating MCCs. (A-C) Confocal pictures of control (A) or CDK7 inhibitors (B, C) treated A6-MCI cells stained for Acetylated-α-tubulin (cilia), Centrin (centrioles) and ZO1 (tight junctions). (D) Graphs displaying the percentage of cells with or without Centrin staining and the percentage of cells with or without cilia. n= 521 (DMSO), 526 (LDC4297) and 809 (YKL-5-124) cells. Data were collected from three independent experiments. Fisher’s Exact Test: p< 2.2e-16 (***) for LDC4297 and YKL-5-124 vs DMSO. (E-G) Confocal pictures of epidermis from control (E) or CDK7 inhibitors (F, G) treated *Xenopus* embryos stained for Acetylated-α-tubulin (cilia), Centrin (centrioles) and ZO1 (tight junctions). (H) Graphs displaying the percentage of cells presenting centrioles defect the percentage of cells with cilia defect. n= 341 (DMSO), 337 (LDC4297) and 368 (YKL-5-124) cells. Data were collected from three independent experiments. Fisher’s Exact Test: p< 2.2e-16 (***) for LDC4297 and YKL-5-124 vs DMSO. (I-K) Confocal pictures of control (I) or CDK7 inhibitors (J, K) treated hAEC stained for Acetylated-α-tubulin (cilia), Centrin (centrioles) and ZO1 (tight junctions). (L) Graphs displaying the percentage of cells with or without Centrin staining and the percentage of cells with or without cilia. n= 1551 (DMSO), 732 (LDC4297) and 578 (YKL-5-124) cells. Data were collected from two independent experiments. Fisher’s Exact Test: p< 2.2e-16 (***) for LDC4297 and YKL-5-124 vs DMSO. Scale bar: 15μm (A-C; E-G; I-K).

## Discussion

In this study, we report the generation of the A6-MCI cell line that can be induced in a controlled manner to become MCCs. We show that this cell line recapitulates the main characteristics of MCCs observed *in vivo*. Using this cell line, we characterized *Xenopus* deuterosomes and provide an unprecedented description of their molecular organization. We also assembled the dynamic proteome of MCCs, a unique resource that will help to identify key regulators and effectors of MCC biogenesis. As a proof of principle, we demonstrate that the uncharacterized kinase CDK7 is of critical importance for MCC differentiation in *Xenopus* and human.

Comparison between MCCs of the *Xenopus* epidermis and A6-MCI cell line showed that both amplify their centrioles via deuterosomes that have comparable architectures. It also highlighted differences with deuterosomes of other MCC subtypes. In mouse ependymal cells, deuterosomes appear as individual spheres of uniform size bearing typically 15 procentrioles, whereas in mouse tracheal MCCs, deuterosomes appear as individual spheres of variable size bearing a variable number of procentrioles. Here, an in-depth analysis using expansion microscopy, TEM and tomography reveals that deuterosomal platforms in *Xenopus* MCCs consist of individual spheres loaded with 2-3 procentrioles. The major and striking difference with other types of deuterosomal structures is that individual spheres are most often organized in chains of variable length, which can intersect and form larger globular structures. It is important to stress that those observations were made with a specific antibody against *Xenopus* Deup1, which ensured that we detected native deuterosomes. In contrast, approaches based on fusion protein expression may be difficult to interpret, due to uncontrolled self-aggregation, which has been reported for Deup1 (Yamamoto et al., 2021). Our observations match those of Steinman on *Xenopus* tracheal and epidermal MCCs, made by TEM over 50 years ago, who reported a linear organization of the core deuterosomal material, that he then called the “procentriole organizer” (Steinman 1969). A first intriguing question emerging from these observations is what causes deuterosomes to adopt different forms? A possible explanation would be that variation in the molecular composition of deuterosomes between different MCC subtypes and/or species, could account for variable organizations. The availability of the A6-MCI resource opens the possibility to decipher the *Xenopus* deuterosomal proteome, thus providing a reference dataset to start answering this question. A second open question relates to the possible link between architecture and function of deuterosomes. It is clear that MCC subtypes present different characteristics, regarding both the number of cilia produced and their spatial organization (Boutin and Kodjabachian, 2019; Mahjoub et al., 2022). In the future, it would be interesting to investigate whether deuterosome organization influences these parameters.

During MCC differentiation, MCI acts upstream of a gene regulatory network that controls the exit from the cell cycle and the amplification of centrioles (Ma et al., 2014; Stubbs et al., 2012). It also activates FoxJ1, which is responsible for the transcriptional activation of the motile ciliogenesis program (Thomas et al., 2010). MCI is therefore a master regulator, which proved sufficient to induce MCC differentiation from heterologous cell types in the *Xenopus* epidermis (Stubbs et al., 2012). However, not all cells react to MCI expression in the same way. Here, we have shown that RPE1 and NIH3T3 cells do not initiate differentiation following MCI expression. Similarly, expression of MCI alone in mouse embryonic fibroblasts only induces centriole overduplication, and co-expression with a constitutively active form of E2F4 is required to activate massive centriole amplification and ciliogenesis (Kim et al., 2018). Thus, it is likely that the identity and/or the epigenetic state pre-existing to forced MCI expression impact the capacity for trans-differentiation into MCC.

The A6-MCI culture model is unique and provide significant advantages over other *in vitro* models such as primary culture of mouse ependymal or tracheal cells. In these models, the differentiation of MCC usually takes several days to weeks and requires specific approaches to remove cells that are not of the MCC lineage, or to induce differentiation with air-liquid interface. These characteristics are not suitable for large-scale production of biological samples needed for in depth proteomic approaches. In contrast, the A6-MCI model is highly suitable for these approaches: the culture develops rapidly, is fairly homogeneous, easy to handle and to scale up, since it can be grown in simple culture flasks or dishes. These assets have enabled us to produce the quantity of material required for the identification of the global proteome over MCC differentiation. The proteome dataset reflects the synchronicity and homogeneity of the A6-MCI culture across time, which represents an asset compared to the *Xenopus* embryo, for which proteomic experiments are global and may not allow to analyse rare or modestly abundant cell types, like MCCs that account for 10% only of the total cell pool in the developing epidermis (Peshkin et al., 2019, (Walentek, 2018).

The analysis of the MCC proteome allowed the identification of several kinases, which may fuel investigations into the MCC alternative cell cycle, a fundamental concept that has recently emerged in the field (Al Jord et al., 2017; Choksi et al., 2024; Serizay et al., 2025). A first interesting candidate for future functional studies is AurkA. AurkA and PLK1 cooperate to regulate different aspects of the mitotic cycle, including centriole maturation and their disjunction in G2 (Joukov and De Nicolo, 2018). In MCCs, we have implicated PLK1 in the disengagement of neo-synthetized centrioles from deuterosomes (Revinski et al., 2018). Interestingly, both PLK1 and AurkA levels were found to increase between 16h and 24 h, before decreasing between 24 and 48h, suggesting a potential cooperation in terminating the phase of centriole biogenesis. It will be interesting to further probe the implication of AurkA in this context. Ten Cdks were detected in the A6-MCI proteome. Some are dynamically expressed and others are stable during differentiation. Among them, we further tested the involvement of Cdk7 and showed that inhibition of this kinase at the time of induction prevents the differentiation of A6-MCI cells into MCCs. During the canonical cell cycle, CDK7 plays an instrumental role by phosphorylating CDK1, CDK2 and CDK4/6 (Fisher, 2005). As the latter have all been implicated in the alternative cycle of MCCs (Al Jord et al., 2017; Choksi et al., 2024; Vladar et al., 2018), it is tempting to speculate that they may also be regulated by CDK7 in MCCs. The A6-MCI model is well suited for phosphoproteomic approaches and should allow progress on this front. In addition, beyond its role as a regulator of mitosis phases, CDK7 belongs to the family of so-called transcriptional CDKs involved in the regulation of RNA transcription (Fisher, 2005). As such, it would be interesting to investigate possible variations in transcription in the presence or absence of CDK7 activity. As Cdk7 is the first kinase of this type to be involved in the molecular regulation of MCCs, it represents an attractive paradigm to investigate in more details how transcriptional and post-translational regulation cooperate to control MCC differentiation.

Broadening our analysis of CDK7, we have shown that its role is not limited to A6-MCI but is conserved in the MCCs of *Xenopus* epidermis, as well as in hAEC. We thus demonstrate that the A6-MCI model has a strong predictive value for understanding the mechanisms at play *in vivo* and in human MCCs. As such, it may help identifying factors relevant in the context of ciliopathies. Of note, CDK7 is a promising target in cancer treatment and several specific inhibitors, including those used in the present study, have entered the clinical development phase (Sava et al., 2020). Here, we show that CDK7 is involved in the development of MCCs, which in humans is a lifelong process that could potentially be affected by CDK7 inhibitor treatments. Our model therefore has implications in cancerology since it could reveal potential side effects of anti-cancer therapies.

## Figure legends

**Supplementary figure 1.**
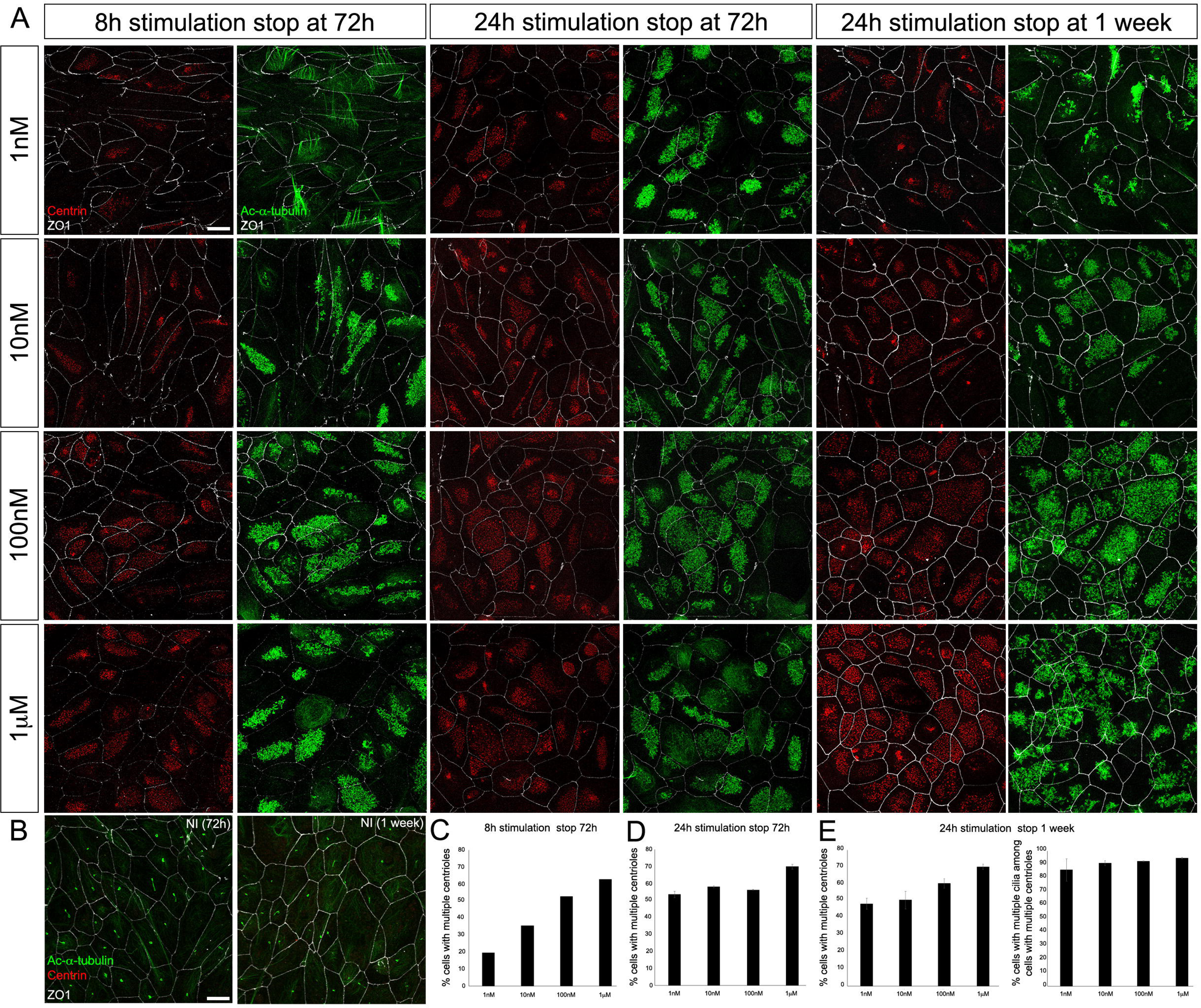
(A) Confocal pictures of A6-MCI induced with indicated concentration of dexamethasone according to indicated protocol and stained for Centrin (centrioles), Acetylated-α-tubulin (Cilia) and ZO1 (Tight junctions). (B) Confocal pictures of non-induced A6-MCI cells stained for Centrin (centrioles), Acetylated-α-tubulin (Cilia) and ZO1 (Tight junctions) after 72h or 1 week of culture. (C) Graph showing the percentage of cells with multiple centrioles at 72hpi for 8h stimulation with increasing dexamethasone concentration. (D) Graph showing the percentage of cells with multiple centrioles at 72hpi for 24h stimulation with increasing dexamethasone concentration. n=936 (1nM), 1277 (10nM), 1432 (100nM) and 1282 (1μM) cells. (E) Graphs showing the percentage of cells with multiple centrioles and with multiple cilia at one week post induction for 24h stimulation with increasing dexamethasone concentration. n=1111 (1nM), 1253 (10nM), 1483 (100nM) and 1379 (1μM) cells. Data were collected from three independent experiments.

**Supplementary figure 2.**
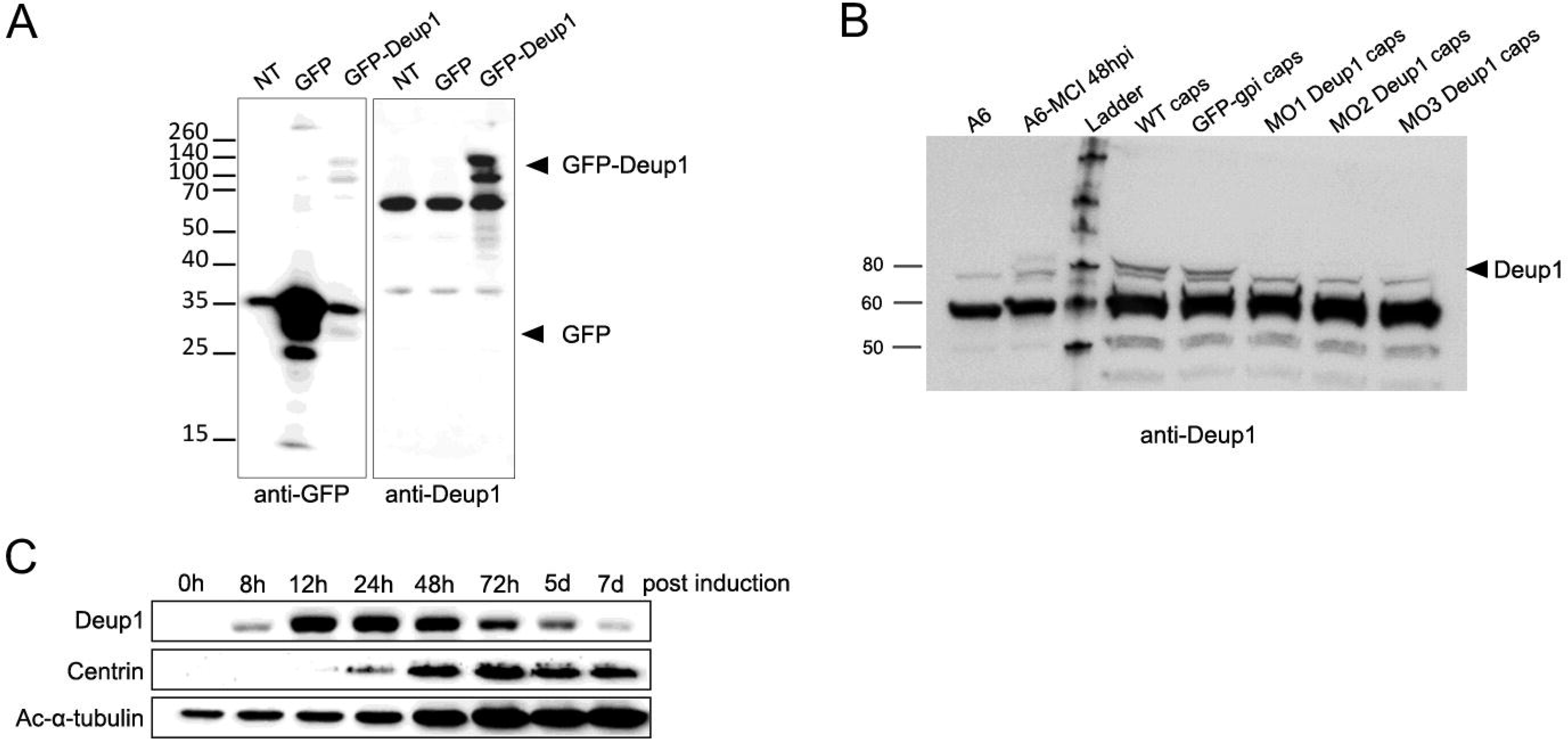
(A) Cos1 cells were transfected with the indicated construct and immunoblotted with anti-GFP (left panel) or home-made anti-Deup1 antibody (right panel). (B) A6, A6-MCI or St18 Xenopus caps injected with indicated construct or MOs were immunoblotted with anti-Deup1 antibody. Deup1 specific band disappear in Caps injected with 3 distinct morpholinos against Deup1. (C) Western blot showing the expression of Deup1, Centrin and Acetylated-α-tubulin in A6-MCI cells at indicated time post induction.

**Supplementary figure 3.**
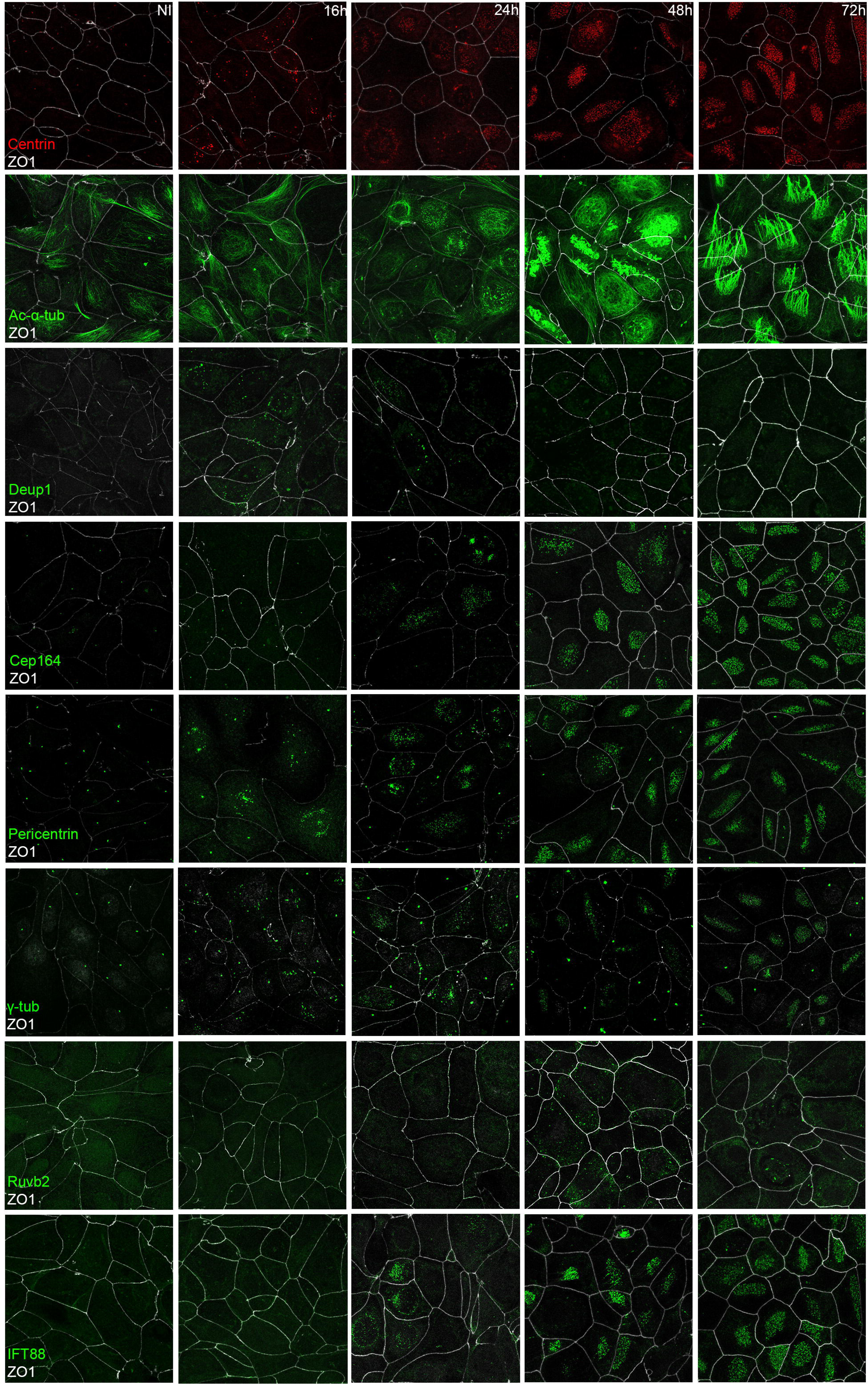
Confocal pictures of A6-MCI cells at indicated time post induction stained with indicated markers.

**Supplementary figure 4.**
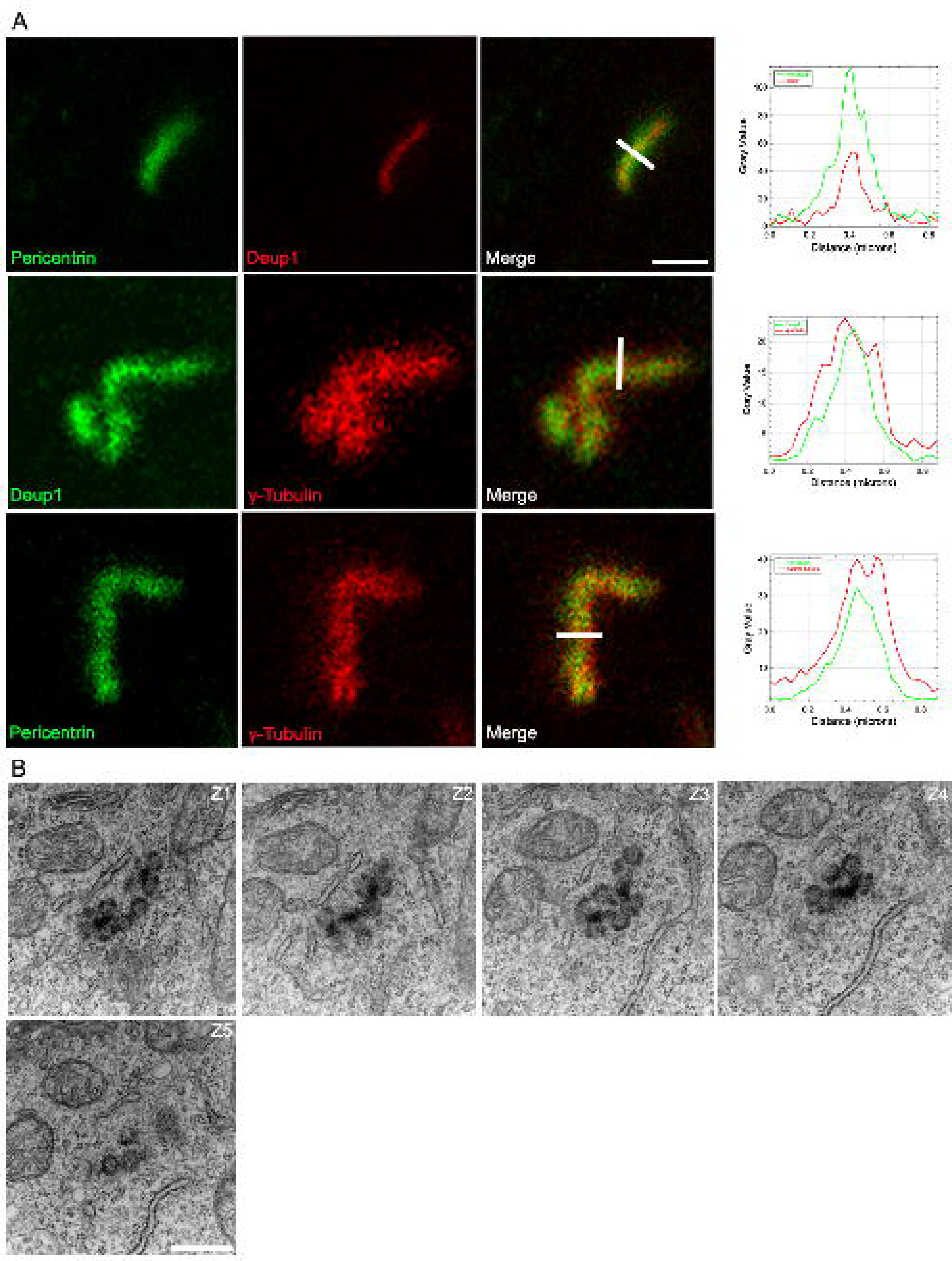
(A) STED pictures of A6-MCI deuterosome revealed by indicated markers. Graphs on right side plot mean intensity grey value for indicated labelling revealing the relative distribution of the proteins. (B) TEM pictures of consecutive 70nm sections through deuterosome of A6-MCI cell. Scale bar: 100nm (A); 500nm (B, C).

**Supplementary figure 5.**
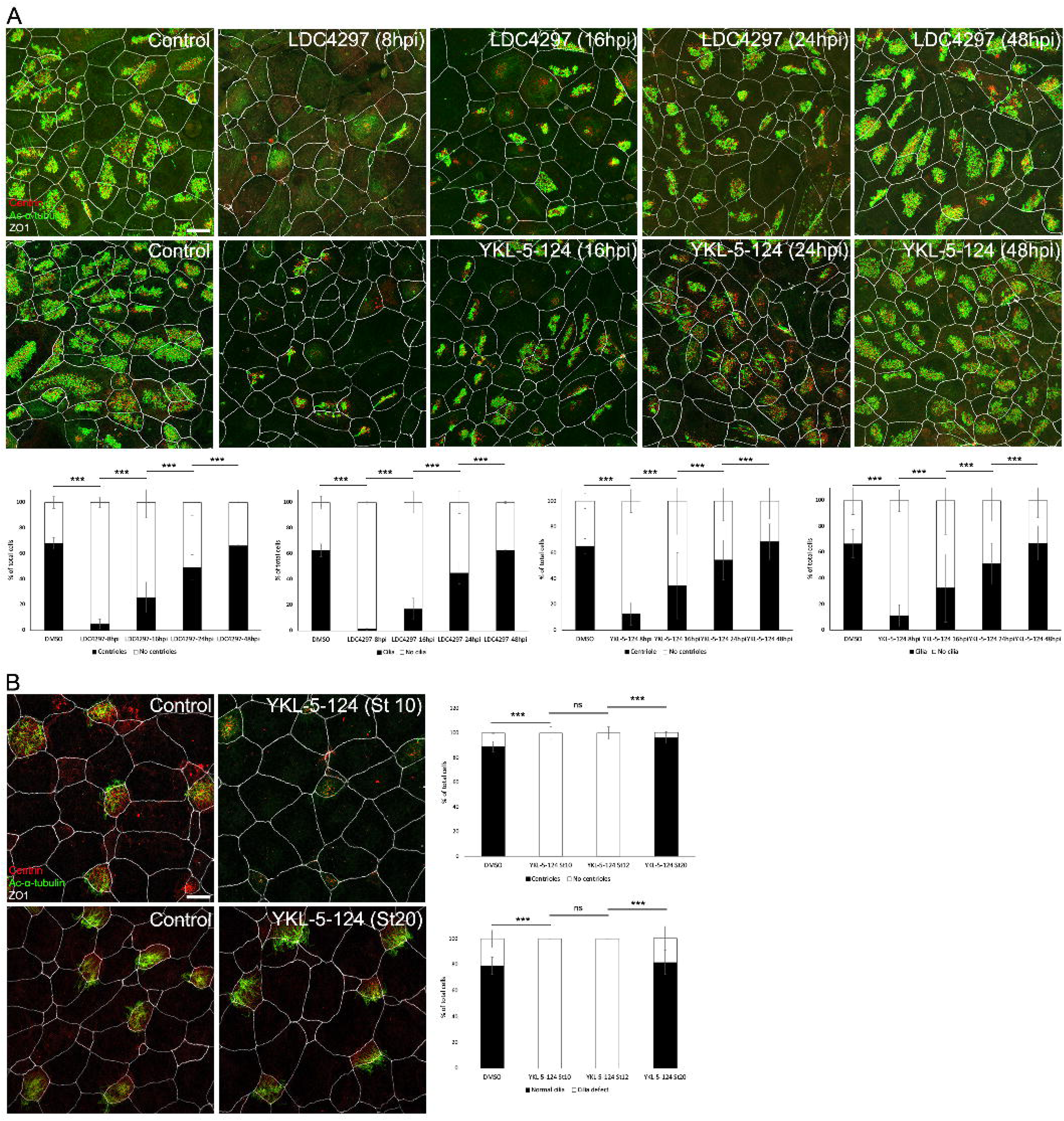
(A) Confocal pictures of A6-MCI cells treated with DMSO (control), LDC4297 or YKL-5-124 at indicated hpi and immunostained for Centrin (Centrioles), Acetylated-α-Tubulin (Cilia) and ZO1 (Tight Junctions) at 72hpi. Graph show quantification of the phenotype. DMSO n= 508, LDC4297 n= 452 (8hpi), 492 (16hpi), 530 (24hpi), 502 (48hpi) and YKL-5-124 n= 494 (8hpi), 527 (16hpi), 526 (24hpi), 622 (48hpi), cells. Data were collected from three independent experiments. Fisher’s Exact Test: p< 2.2e-16 (***) for all comparisons. (B) Confocal pictures of *Xenopus* epidermis cells treated with DMSO (control) or YKL-5-124 at indicated embryonic stages and immunostained for Centrin (Centrioles), Acetylated-α-Tubulin (Cilia) and ZO1 (Tight Junctions) at stage 27. Graph show quantification of the phenotype. DMSO n= 125 (DMSO) and YKL-5-124 n= 124 (St10), 141 (St12), 160 (St20) cells. Data were collected from three independent experiments. Fisher’s Exact Test: p< 2.2e-16 (***) for DMSO vs YKl-5-124 (St10), YKl-5-124 (St12) vs YKl-5-124 (St20); p=1 (ns) for YKl-5-124 (St10) vs YKl-5-124 (St12).

## Supplementary movie 1

Tomogram acquisition and 3D tomogram reconstruction of short deuterosome.

## Supplementary movie 2

Tomogram acquisition and 3D tomogram reconstruction long deuterosomes.

## Supplementary movie 3

Tomogram acquisition and 3D tomogram reconstruction of a ramified deuterosomes. Deuterosome core is shown in green, centrioles are in red and undefined material in white.

## Supplementary table 1

Lists of proteins differentially expressed by A6-MCI between post-induction time points. Up-regulated proteins with a Log2 difference >0.5 (Student’s T-test q-value < 0,05 and Log Student’s T-test p-value <1.3) and down-regulated proteins with a Log2 difference <-0.5 (Student’s T-test q-value < 0,05 and Log Student’s T-test p-value <1.3) are displayed.

## Material and methods

### Cells and transfection

A6 xenopus cells were obtained from the American Type Culture Collection and grown at 27°C in complete medium composed of 55% Leibovitz’s L15 medium, 10% heat inactivated fetal bovine serum (FBS), 20 u/ml penicillin, 20 µg/ml streptomycin (Thermo Fisher Bioscience). Cells were transfected with Fugene HD (Promega) according to the instructions of the manufacturer. Cells were grown in the presence of 1.5 mg/ml G418 for selection and subsequently 0.5 mg/ml for maintenance.

### Plasmids

A pCS2+ plasmid coding for Xenopus laevis MCIDAS fused to the ligand binding domain of the human glucocorticoid receptor (Stubbs et al., 2012) was obtained from Chris Kintner and modified by the insertion of a loxP flanked Neor cassette (Arakawa et al., 2001) at NotI/KpnI sites. An expression vector coding for Myc-tagged mouse MCIDAS (plentiPGK-Myc-Mcidas) was obtained from Eszter Vladar.

### Induction of A6-MCI differentiation and drug treatment

10^6^ cells were seeded per well of six wells culture plate and grown at 27°C to confluency in complete G418 medium. The medium was replaced for 24 h with 1% FBS containing medium without G418 and subsequently supplemented with 100nM (or concentration specified in figures) dexamethasone for additional 24 h or less. Depending on the total incubation time (>24 h) the medium was changed to the same medium without dexamethasone before fixation or lysis. YKL-5-124 and LDC4297, diluted in DMSO at 10μM and 2.5μM respectively were added to culture medium either at time of induction or at different time post-induction. Control cells were treated with 0.1% DMSO.

### *Xenopus* embryos culture, drug treatment and morpholinos injections

Eggs obtained from NASCO females were fertilized in vitro, dejellied and cultured using standard protocols (Revinski et al., 2018). At stage 12, vitelline membranes of the embryos were removed and 400 µM of LDC4297 and 200 µM of YKL-5-124 diluted in DMSO were added to the culture medium. Control embryos were treated with 0.4% DMSO. Embryos were incubated at 18°C until they reach stage 27-30. Deup1 translation (MO1: 5’-GGCTTTCAGTGTCTGTTTGCATTTC-3’ (Mercey et al., 2019) ; MO2: 5’-TGTGTCTCCGGCTCCCAGATAAAAC-3’) or splice (MO3: 5’-AAGGAAACAAACCACACTCACCTAC-3’) blocking morpholinos were injected in the 4 blastomeres at NF stage 3. Animal caps from WT embryos or from embryos injected with GFP-GPI (control) or Deup1 MO were obtained by manual dissection from stage 10 (n=50 embryos/condition) in 1× MBS and kept in 0.5× MBS until matched control embryos reached stage 18.

### human Airway Epithelial Cells culture and drug treatment

Fresh cryopreserved Human Bronchial Epithelial cells (HBEpC, C-12640, PromoCell, Max Passage 3) were thawed in a T75 in Complete PneumaCult-Ex Plus medium and grown in an incubator at 37°C with 5% CO2. A medium change was performed 24h after defrosting and every two days until cells reach 50-60% confluence. After trypsination 250 000 cells/ml were seeded in Transwells (Corning 6.5 mm Transwell with 0.4 µm Pore polyester Membrane Insert, n°3470, Costar) in Complete PneumaCult-Ex Plus medium apical and basal chambers. Just before confluence, cells were placed in air-liquid interface (ALI) by removing medium from the apical chamber and replacing medium in the basal chamber by complete PneumaCult-ALI medium supplemented with 0.25% DMSO (control) or 50 nM LDC4297 or 25 nM YKL-514 diluted in DMSO. Medium change was performed every two days and drugs treatments were renewed at the same time. From ALI day 7, dPBS 1X apical washes were performed twice a week to remove excessive mucus. From ALI day 23, differentiation of cells was monitored daily using a Nikon eclipse Ti inverted microscope in brightfield mode. When DMSO controls reached 50% differentiation cells were fixed in cold methanol at -20°C during 6 min and proceeded for immunostaining.

### Cell lysis and Western blotting

Cells were washed in PBS and lysed in 50 mM Tris-HCl pH 7.5, 150 mM NaCl, 1mM EDTA, containing 1% NP-40 and 0.25% sodium deoxycholate (modified RIPA) plus a Complete Protease Inhibitor Cocktail (Roche Applied Science) on ice. Cell extracts separated on polyacrylamide gels were transfered onto Optitran membrane (Whatman) followed by incubation with primary antibodies and horseradish peroxidase conjugated secondary antibodies (Jackson Immunoresearch Laboratories). Signal obtained from enhanced chemiluminescence (Western Lightning ECL Pro, Perkin Elmer) was detected with MyECL Imager (Thermo).

Animal caps were lysed in 200 µl of RIPA buffer (50 mM Tris-HCl pH 7.5, 150 mM NaCl, 1% NP40, 0.1% SDS and 0.5% sodium deoxycholate) containing a protease inhibitors tablet (Pierce). Total protein concentration was determined by Bradford assay (Invitrogen) and samples were prepared in LDS sample buffer (Invitrogen) containing a reducing agent. Samples in LDS buffer were denatured for 5 min at 100°C and 120 µg of proteins were loaded and separated on 4–20% SDS-PAGE (Mini-PROTEAN® TGX™ #456109, Bio-Rad). Following migration, proteins were transferred onto 0.45 µm nitrocelullose membranes (GE Healthcare). Membranes were blocked in Tris-buffered saline (TBS) for 1 h at RT and then incubated overnight at 4°C with the primary antibody. After three washes in TBS with 0.05% Tween 20 (TBST) buffer for 10 min, the membrane was incubated with the appropriate HRP-conjugated secondary antibody (Jackson ImmunoResearch) diluted at 1:5000 in TBST-5% dry fat milk and signals were detected using chemiluminescence.

### Generation of Xenopus Deup1 antibody

Homemade rabbit antibodies were produced by immunization with recombinant portion of *Xenopus laevis* DEUP1 (XP_018103272.1, residues 1-280).

### Immunofluorescence staining

A6-MC cells were grown on glass coverslips and fixed for 6 min in methanol at -20°C. Cells were washed in PBS, blocked in PBS, 3% BSA and stained with primary antibodies (see table below) in blocking buffer. After washings in PBS 0.1% Tween-20, cells were incubated with fluorophore conjugated secondary antibodies (see table below), washed and DNA was stained with 250 ng/ml DAPI. Coverslip were then rinsed and mounted in Mowiol.

*Xenopus* Embryos were fixed at RT in PFA 4% - 0.1% Triton X100. Embryos washed in PBS, blocked in PBS-BSA 3% and incubated overnight with primary antibodies. After washing in PBS, embryos were incubated 1 hour at RT with fluorophore conjugated secondary antibodies (see table below) at RT. Embryos were washed with PBS 1X, before mounting with Mowiol between slides and coverslips.

hAEC fixed with methanol were washed 3 times with PBS 1X – Tween 20 0.1% before blocking in PBS-BSA 3% 15min at RT. Cells were incubated 45min at RT with primary antibodies (see table below) diluted in PBS 1X -BSA 3%. After 2 washes with PBS 1X – Tween 20 0.1% cells were incubated 30min at RT with secondary antibodies (see table below) diluted in PBS 1X-BSA 3% at RT. After 2 washes with PBS 1X – Tween 20 0.1%. Transwell membranes were cut before mounting with Mowiol between slides and coverslips.

### Expansion microscopy

A6-MCI cells were grown on glass coverslips and fixed 6min in MetOH before U-ExM processing according to published protocols (Gambarotto et al., 2019). Briefly, cells were incubated for 5h in 1.4% formaldehyde (FA) / 2% acrylamide (AA) at 37°C. FA/AA solution was removed and replaced by gelation solution composed of monomer solution containing 19% (wt/wt) SA, 10% (wt/wt) AA, 0.1% (wt/wt) BIS in 1× PBS supplemented with 0.5% APS and 0.5% TEMED. Gelation step proceed 5min on ice and 1h at 37°C. Coverslips with gel were transferred in 6 wells plate with denaturation buffer (200 mM SDS, 200 mM NaCl, and 50 mM Tris in ultrapure water, pH 9) for 15min at RT. Gel were transferred to fresh denaturation buffer in 1.5ml Eppendorf and incubated at 95°C for 30min. After denaturation, gels were placed in beakers filled with ddH2O for the first expansion. Water was exchanged at least twice every 30 min at RT, and then gels were incubated overnight in ddH2O. Gels were incubated in PBS1X 2X 15min before blocking in PBS-3%BSA for 3h at 37°C. Primary antibodies (see table below) diluted in PBS-2%BSA were added for 3h at 37°C. After wash with PBST 3X10min gels were incubated with secondary antibodies (see table below) diluted in PBS-3%BSA for 3h at 37°C. Gels were washed 3X 10 min under agitation and place in ddH2O for final expansion. ddH2O was exchanged at least twice every 30 min, and gels were incubated in ddH2O overnight. Gels expanded around 4 time.

### Antibodies

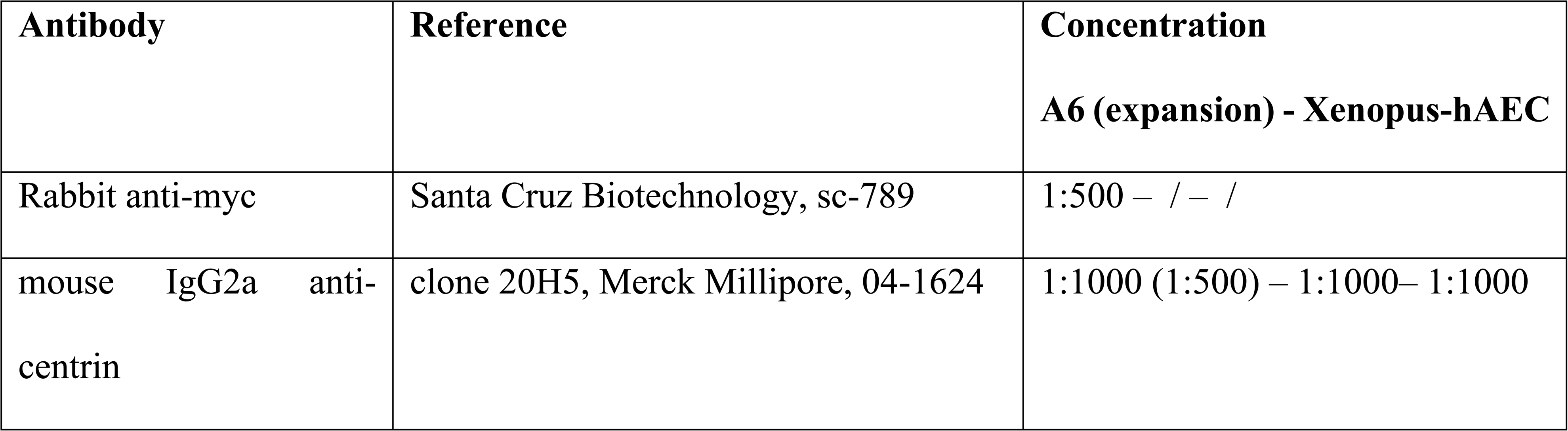
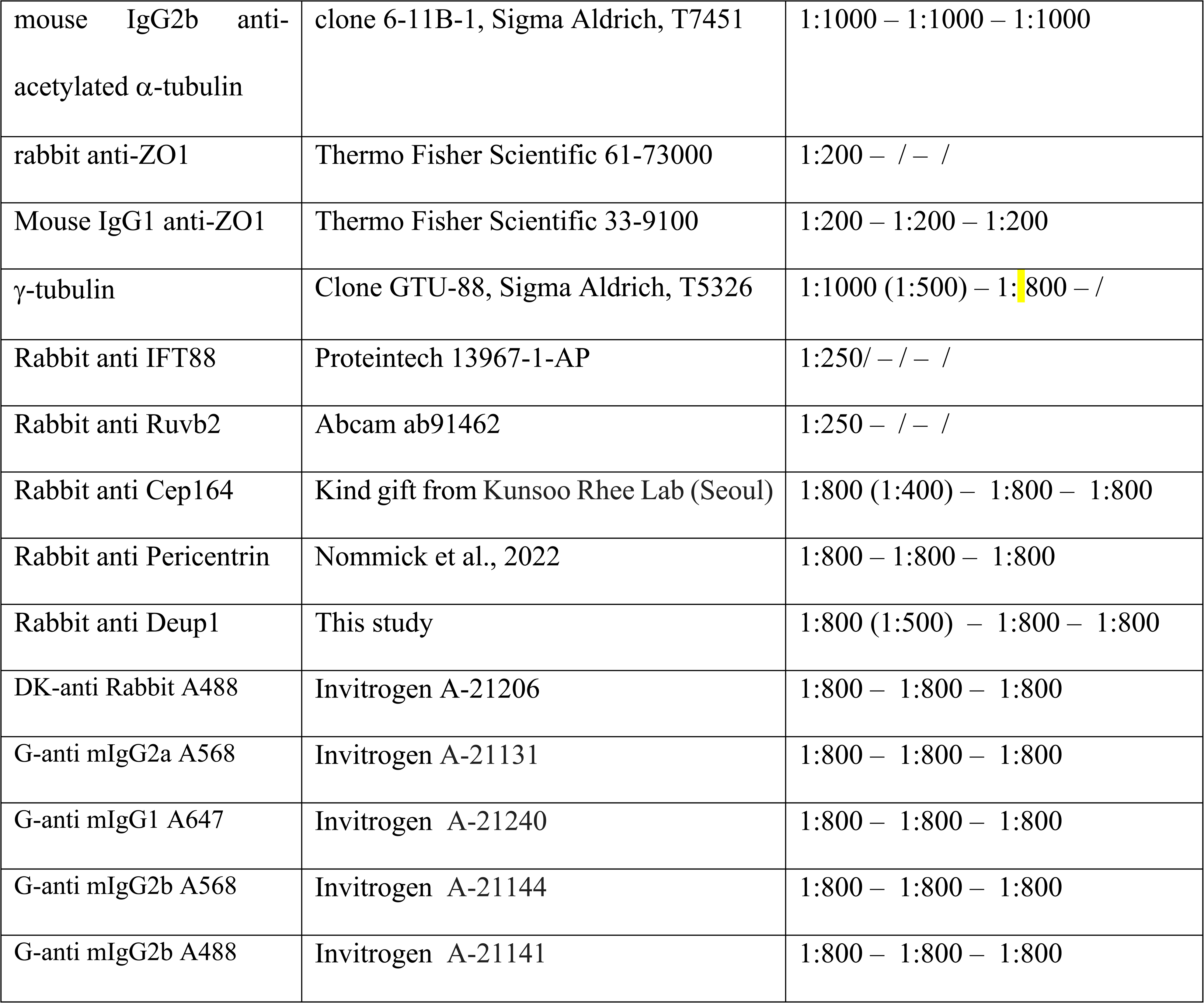

### Pictures acquisition and quantification

Confocal images were acquired by capturing Z-series with 0.3 μm step size using a LSM 780 or 880 (Zeiss) or SP8 (Leica microsystems) laser scanning microscope equipped with 63X Oil objectives. Images were converted into single plane maximum intensity projection (MIP) and processed using Fiji software. STED super-resolution images were acquired with a TCS SP8 STED 3X microscope equipped with an HC PL APO 93X/1.30 GLYC motCORRTM objective (Leica microsystems). CSD quantifications was done using Biotool Software as previously described (Boutin et al., 2014). Prism software or Excel were used for graphical representation. The average maps of Fig.3D were obtained through the following three steps: manual picking of 13 particles in the confocal data (Fig.3C); reference-free reconstruction of a coarse average map using the method described in (Eloy et al., 2023); refinement of this initial map using the approach detailed in (Fortun et al., 2016). All these steps were performed with the publicly available SP-Fluo software1. 1https://spfluo.icube.unistra.fr/en/usage/installation.html

### Transmission electron microscopy, tomography and segmentation

Cells were fixed with glutaraldehyde 2.5 %, paraformaldehyde 2 % in Cacodylate buffer 0.1 M pH 7.4, and post-fixed with 1 % Osmium Tetraoxyde in Cacodylate buffer 0.1 M pH 7.4 during 1 hour at RT. They were dehydrated in alcohol and embedded in Epon. Next, cells were cut transversely (70 nm/slice for serial sections 300nm/slice for tomograms) with a Leica Ultracut UC7 (Leica, Germany) and the serial sections were collected on grids. In order to observe and reconstruct entire deuterosomes, images or tomograms were acquired on several consecutive sections until no deuterosomal material was visible on the edge sections. Images were acquired using a Tecnai G2 (Thermofisher, USA) microscope, running an LaB6 crystal at 200kV and equipped with a 2K Veleta camera (Olympus, Japan). For tomograms, dual-axis tilt series were acquired at 62kx (0.94 nm/px) or 80kx (0.73 nm/px) following the saxton scheme (0° tilt step : 1°) from -60° to +60°.

Tomographic reconstructions were carried out in IMOD, using cross-correlation for tilt series alignment and back projection algorithm for reconstruction. The IMOD MIDAS semi-automated alignment tool was used to join serial tomograms.

The reconstructed volumes were denoised by the anisotropic diffusion filter in Amira and treated with the FeatureJ plugin in Fiji before segmentation by pixel classification in ilastik. Probability maps of segmentation were imported into Dragonfly were the core versus the centriolar material were discriminated using the multi-slice brush tool in adaptive Gaussian mode. Segmented volumes were finally exported to Amira for movie generation

### Proteomic

For each condition, 2X 10^7^ cells were seeded in 10 cm culture dish and grown at 27°C to confluency in complete G418 medium. The medium was replaced for 24 h with 1% FBS containing medium without G418 before induction with 100 nM dexamethasone. Cells were washed twice with cold PBS1X and incubated 4min in RIPA lysis buffer complemented with protease and phosphatase inhibitor (ThermoFisher Scientific, #78441). Cell lysate was collected, incubated 15 min on ice with nuclease (ThermoFisher Scientific, #88700) and centrifuged 25min at 16000g at 4°C. Cell extracts were collected, proteins were measured using a BCA kit, and samples were adjusted to a concentration of 2 mg/ml in RIPA buffer. 15µg of each protein extract was stacked on 4–12% Bis–tris acrylamide gels NuPAGE™ (Life Technologies) in a single band to perform trypsin digestion before mass spectrometry analysis using an Orbitrap Fusion Lumos Tribrid (ThermoFisher Scientific, San Jose, CA) in data-independent acquisition mode (DIA). Protein identification and quantification was processed using the DIA-NN 1.8 algorithm (Demichev et al., 2020) and DIAgui package (https://github.com/marseille-proteomique/DIAgui (Gerault et al., 2024)). See supplementary methods for a detailed method. The mass spectrometry proteomics data have been deposited to the ProteomeXchange Consortium via the PRIDE partner repository with the data set PXD06447 (Perez-Riverol et al., 2025). The statistical analysis was done with the Perseus program (version 1.6.15.0) (Tyanova and Cox, 2018). Differential proteins were detected using a two-sample t test at 0.05 permutation-based false discovery rate and an additional filter on absolute Log2 difference >0.5. Statistical analysis was performed using the standard two-tailed Student t test and p value < 0.05 were considered significant. Volcano plots were created using Prism software. GO analysis was performed using STRING data base (Version 12.0).

### Data availability

Data will be made available on request. Proteomics data have been deposited to the ProteomeXchange Consortium via the PRIDE partner repository with the dataset identifier PXD06447.

## Acknowledgements

This work received support from the French government under the France 2030 investment plan, as part of the Initiative d’Excellence d’Aix-Marseille Université – A*MIDEX” AMX-21-PEP-042 (‘Pépinière d’Excellence 2021 to C.B.), Cancéropôle PACA-GEFLUC (Programme Emergence 2021 to C.B.), ANR (ANR-22-CE13-0027 PHACIL to C.B. and MCCproteome ANR-19-CE13-0033 to L.K.). Imaging was performed at IBDM, member of the National Infrastructure France-BioImaging (https://ror.org/01y7vt929) supported by the French National Research Agency (ANR-24-INBS-0005 FBI BIOGEN). We thank Emmanuelle Leblanc and Florian Roguet for *Xenopus* husbandry. J-P. B. is a scholar of the Institut Universitaire de France. The mass spectrometry facility of Marseille Proteomics (https://marseille-proteomique.univ-amu.fr/) is supported by IBISA, the Cancéropôle PACA, the Provence-Alpes-Côte d’Azur Region, the Institut Paoli-Calmettes, and Fonds Européen de Développement Regional (FEDER). We thank Julia Schaeffer and Peter Walentek for insightful comments on the manuscript.

## Authors contribution

Conceptualization: L.K. and C.B.; Investigation: C.B., O.R, M.T., S.D., N.B, V.T., and D.F.; Proteomics expertise: S.A. and L.C.; Supervision of proteomics: J.P.B.; Formal analysis: C.B.; Writing: C.B.; Editing: all authors.; Funding acquisition: L.K. and C.B.; Supervision: C.B.

## Declaration of interest

The authors declare no competing interest.

## Notes

### Competing Interest Statement

The authors have declared no competing interest.

